# Nuclear Histone 3 Post-Translational Modification Profiling in Whole Cells using Spectral Flow Cytometry

**DOI:** 10.1101/2024.10.03.616268

**Authors:** Carly S. Golden, Saylor Williams, Sandeep Sreerama, Sophia Blankevoort, H. Joseph Yost, Martin Tristani-Firouzi, Anna Belkina, Maria A. Serrano

## Abstract

Histone modifications play essential roles in regulating chromatin accessibility and downstream transcription, serving as critical determinants of cell identity and function. However, the diversity of histone post-translational modifications (PTMs) and their tendency to be studied in isolation limits our understanding of their coordinated roles in shaping cellular states. Conventional flow cytometry methods for histone PTM assessment suffer from low multiplexing capacity and typically require nuclear isolation, resulting in significant sample loss and an incomplete picture of the overall cell state. Here, we present EpiFlow, a spectral flow cytometry protocol designed for multiparametric analysis of histone PTMs within whole cells. By utilizing a 96-well plate format and preserving the cell entirely, EpiFlow improves throughput and efficiency while retaining sample integrity. This method resolves subtle variations in histone PTMs within neural progenitor cells, capturing distinct chromatin states across the cell cycle and correlating them with several markers. Furthermore, we provide an open-access GitHub repository containing detailed protocols and analysis workflows, ensuring reproducibility and accessibility of this approach. EpiFlow offers a robust framework for exploring chromatin dynamics, with broad implications for advancing fundamental research and therapeutic research.

## Introduction

Epigenetic proteins are key regulators of chromatin accessibility and cell differentiation (Allis and Jenuwein, 2016; Chen and Dent, 2014; Jiang et al., 2011; Santos-Rosa et al., 2002; Stergachis et al., 2013). Understanding these processes within the context of cell status is crucial, as cellular states emerge from coordinated changes across molecular events. The application of large scale integrated analytic methods, such as those used in systems biology (Chuang et al., 2010) and omics studies (Joyce and Palsson, 2006), provides a simultaneous view of individual molecular events within the context of broader biological processes. This approach offers deeper insights into how small molecular changes collectively drive cell fate and function. This is especially relevant for epigenetics, where histone modifications govern DNA access that guide differentiation events during development. Indeed, chromosome structure and epigenetic modifications are known to undergo significant changes during germline induction (Weng et al., 2012), cell differentiation (Grigoryev et al., 2006; Larson and Yuan, 2012), maintenance of cell identity (Yiyuan Liu et al., 2017), and throughout the cell cycle (Calder et al., 2013; Coronado et al., 2013; Petruk et al., 2012). Epigenetic modifications have also been used as clinical indicators for assessing and predicting disease outcomes, as discussed in the following reviews (Egger et al., 2004; Kelly et al., 2010; Lim et al., 2024). By employing integrative methods at the cellular level, we can better link histone modifications to broader cellular outcomes with clinical significance.

Histone profiling using flow cytometry enables the study of global histone modification changes, providing detailed insights into chromatin dynamics at the single-cell level (Bergamasco et al., 2024; Ronzoni et al., 2005). However, this approach is currently limited by low multiplexing capability, the need for nuclear isolation, and insufficient resolution to detect subtle biological variations, providing an incomplete view of the broader cellular state. To address these challenges, we developed EpiFlow, a spectral flow cytometry protocol that improves the resolution and throughput of histone post-translational modification (PTM) profiling while simultaneously capturing diverse attributes of cell status. Using human induced pluripotent stem cells (hiPSCs) and their neural progenitor cell (NPC) derivatives, we aimed to capture the interplay between histone PTMs, cell cycle phases, apoptosis, viability, and cell lineage markers as it relates to unique cell states.

Spectral flow cytometry represents a significant advancement over traditional flow cytometry methods, offering unparalleled multiplexing capabilities and higher sensitivity for detecting subtle biological changes (Futamura et al., 2015; Schmutz et al., 2016). Unlike conventional approaches, spectral flow cytometry captures the full emission spectrum of each fluorophore, enabling the resolution of highly complex staining panels (Nolan and Condello, 2013; Novo, 2022). This capability has been leveraged in recent studies to uncover intricate relationships between cell identity and function, making it a powerful tool for both basic research and clinical applications. For example, spectral flow cytometry has been used to identify rare cell populations (Vanuytsel et al., 2022) and resolve cellular processes with clinical relevance, such as immune profiling in cancer (DeNiro et al., 2024; Sanjabi and Lear, 2021; Spasic et al., 2024; Valenzano et al., 2024). By combining these advantages with a robust experimental workflow, spectral flow cytometry provides deeper biological insights that were previously unattainable.

Building on these advancements, we significantly optimized the application of spectral flow cytometry for epigenetic research. EpiFlow further addresses limitations of traditional histone profiling methods by employing a whole-cell staining protocol in a high-throughput 96-well plate format. This eliminates the need for nuclear isolation, minimizing sample loss and enabling simultaneous detection of membrane-bound, cytoplasmic, and nuclear antigens. Additionally, our protocol integrates multiplexing capabilities to profile multiple histone PTMs, apoptotic and proliferating cells, and lineage-specific transcription factors in a single assay. By leveraging high-dimensional data readouts that concurrently considers fluorescent signals from all fluorophores within a single cell—a distinctive feature unique to spectral flow cytometry—we achieve greater resolution and sensitivity that enables us to answer more complex questions about chromatin biology. Finally, by streamlining data acquisition and analysis through our open-access GitHub repository (Golden et al., 2024; GitHub Repository), we further ensure reproducibility and accessibility for the broader scientific community (Ushey and Wickham, 2024a, 2024b).

Through the application of EpiFlow, we identified dynamic changes in histone PTMs and lineage-specific markers across the cell cycle in hiPSC-derived NPCs. By enabling simultaneous profiling of histone PTMs and cell status markers, EpiFlow provides a higher-resolution view of epigenetic regulation than previously possible. These results highlight the utility of EpiFlow as a versatile tool for exploring epigenetic regulation and cell differentiation, with implications for understanding developmental biology and advancing therapeutic strategies. Ultimately, this work underscores the potential of spectral flow cytometry to bridge the gap between single-cell analyses and systems-level insights into chromatin dynamics.

## Results

### Antibody Panel Design for Multiparametric H3-PTM Profiling and Cellular Features

Spectral flow cytometry employs more sensitive detectors that capture the full emission spectrum of each fluorochrome, permitting the use of larger panels to evaluate broader cell attributes (Ferrer-Font et al., 2021). Unique spectral signatures of each fluorochrome are identified through advanced unmixing techniques, addressing the compensation limitations observed in traditional flow (Novo, 2022). However, the limited availability of conjugated and validated monoclonal antibodies for histone PTMs represents a challenge for multiparametric panel design. To address this limitation during the design and validation of the EpiFlow panel, we selected histone PTM antibodies that met the following criteria: (a) availability of monoclonal antibodies conjugated with fluorochromes that have compatible spectral signatures for multiplexing, (b) histone marks with published literature supporting their association with transcriptional activation and distinct cell cycle phases; H3K27ac (Yin Liu et al., 2017; Yiyuan Liu et al., 2017), H3K4me3 (Lauberth et al., 2013), H3K9ac (Krejčí et al., 2009), H3K14ac (Kueh et al., 2023), H3K14ac+pH3 (Lo et al., 2000), H3K9ac+H3K14ac (Armenta-Castro et al., 2020; Karmodiya et al., 2012; Liu et al., 2023) and, (c) known responses to small molecules that will increase or decrease histone mark levels for validation purposes. This resulted in an initial histone H3 PTM core panel for H3K9ac, H3K14ac, H3K4me1, H3K4me3 and H3K27ac. Additional features to identify live/dead cells, total H3, distinct cell types, mitotic cells, apoptotic cells, and to resolve cell cycle phases were added based on spectral compatibility with the core H3-PTM panel (Table S1). Our staining protocol was optimized to measure nuclear histone PTMs and features of interest across subcellular compartments in fixed whole cells. However, a significant challenge of this multiplexing approach is maintaining the integrity of membrane-bound epitopes while ensuring optimal fixation and permeabilization to allow antibodies sufficient access to the nuclear compartment and effective antigen recognition. To validate our staining protocol for cell membrane integrity and nuclear antigens accessibility, we performed immunofluorescence followed by confocal imaging of SCTi-003A hiPSCs stained with H3K9ac and beta-catenin (a membrane marker) (Fig. S1). Our results demonstrate proper staining and the expected expression patterns across cell compartments, validating this protocol for compatible staining of histone PTMs, nuclear, and membrane-bound epitopes. This ensures reliable multiplexed detection for downstream spectral flow cytometry assays.

### Validation of EpiFlow for Sensitive and Specific Detection of Histone H3 Acetylation and Methylation Marks

To assess the specificity of the selected antibodies, we treated SCTi-003A hiPSCs and WTC11 hiPSC-derived NPCs and with Trichostatin-A (TSA), SAHA (Vorinostat), or A-485 to selectively modulate histone H3 acetylation levels (Lasko et al., 2017; Richon et al., 1996; Seuter et al., 2013). TSA and SAHA are histone deacetylase I/II inhibitors (HDACi) that increase histone acetylation, while A485 is a histone acetyltransferase inhibitor (HATi) targeting the HAT coactivator complex p300/CBP, which ultimately reduces histone acetylation. We evaluated EpiFlow sensitivity using three different concentrations of these selective inhibitors and tested whether we could detect subtle changes in H3-PTMs across cell cycle phases. Our experiments in hiPSCs and NPCs showed a significant and dose-dependent change, either increasing (TSA and SAHA groups) or decreasing (A-485 group) median fluorescent intensity values (MFI) of H3K27ac, H3K4ac, and H3K9ac marks (Fig. S2, S3). Our results also showed distinct changes in the distribution of MFI values of each H3-PTM depending on both the expression levels of PAX6 (NPC marker) and cell cycle phase (Fig. S2 D, E). MFI is one of the most common statistics used to measure the shift in fluorescence intensity (FI) of a cell population. However, MFI values report the central tendency of a cell population and do not provide information about the overall fluorescent distribution for a feature of interest (Herzenberg et al., 2006; Roederer et al., 2004). This limitation is particularly relevant for experimental designs aimed at detecting subtle changes in a feature’s fluorescent profile, such as histone H3 acetylation and methylation levels, as a response to drug treatments. TSA and A-485 are highly specific HDAC and HAT inhibitors, respectively, widely used to modulate histone H3 acetylation levels (Lasko et al., 2017; Vigushin et al., 2001). Therefore, we proposed that TSA and A-485 treatments would not change the FI values of histone H3 methylation marks (H3K4me), further validating EpiFlow specificity. Interestingly, H3K4me1 FI values showed significant differences between drug treatments and across cell cycle phases (Krustal-Wallis test G0/G1 χ^2^(6) = 4380.63, *p*<0.0001; G2 χ^2^(6) = 817.47, *p*<0.0001; Mitosis χ^2^(6) = 236.45,

*p*<0.0001, Supplementary GitHub analysis, Methods Validation section). Further, the TSA group showed an overall more pronounced effect on increased H3K4me1 signal than the A-485 group, particularly in mitosis. Despite this, for all treatment groups, H3K4me1 MFI values shifted in the same direction towards higher intensity values without affecting the overall signal distribution (Supplementary GitHub analysis, Methods Validation section). We believe the observed H3K4me1 changes could indicate a technical limitation of our approach or suggest an alternative scenario in which modulation of H3 acetylation might also influence H3K4 methylation levels (Bui et al., 2010; Crump et al., 2011; Lee et al., 2006). This prompted us to validate the sensitivity of our panel with two independent approaches: conventional flow cytometry to measure H3K4 methylation levels in hiPSCs treated with either OICR-9429 or CPI-455 HCl, and in an hiPSC line with a homozygous loss of function (LOF) mutation in the histone methyltransferase gene *KMT2D* (KMT2D10.4). OICR-9429 is a potent antagonist that inhibits the WDR5-MLL complex responsible for mediating H3K4me3, consequently reducing this mark (Grebien et al., 2015). CPI-455 HCL selectively inhibits the lysine demethylase KDM5, resulting in increased global H3K4me3 levels (Vinogradova et al., 2016). KMT2D10.4 hiPSCs were generated using a CRISPR/Cas9-targeted 2 base pair deletion in exon 2 of *KMT2D* gene in a WTC11 hiPSC line (File S1). The resulting line is expected to exhibit reduced levels of H3K4 methylation compared to its isogenic control. WTC11 hiPSCs treated with OICR-9429 and CPI-455 HCL showed the expected shift in both the magnitude and direction of H3K4me3 FI (Fig. S4 C, E). This was further supported by a significant reduction of H3K4me2 and H3K4me3 FI values in the *KMT2D* LOF line compared to its isogenic control (Fig. S4 F). Although still significant, H3K4me1 levels remained relatively consistent between the KMT2D10.4 and WTC11 cells (Fig. S4 A-C), which aligns with recent studies reporting redundant H3K4me1 activity by other KMT2 methyltransferases in mouse embryonic stem cells (Kubo et al., 2024). Our data, generated from three different hiPSC lines (SCTi-003A, WTC11, and KMT2D10.4), in two cell types (hiPSC and NPC), acquired across two instruments (Aurora Cytek and Stratedigm S1000EXi), demonstrate that EpiFlow can detect histone H3 PTMs with high sensitivity and specificity. Additionally, our analysis underscores the importance of resolving the cell cycle status of cell populations to identify histone PTM changes more accurately.

### EpiFlow Application for Resolving H3-PTM Dynamics Across Cell Cycle Phases in NPCs

The simultaneous assessment of several cellular parameters in one experiment allows for high-resolution profiling of complex samples, rare cell populations, or highly dynamic biological processes in specific cell lineages. We hypothesized that EpiFlow could resolve distinct alterations of histone H3 PTM across cell cycle phases in a single cell lineage, and that subtle variations in these modifications would contribute to subcluster formation within each cell cycle phase. We again applied the core EpiFlow panel (H3K4me3, H3K9ac, H3K27ac, H3K14ac and H3S10ph) to hiPSC-derived NPCs and included features that identify live cells (Live/Dead staining), apoptotic cells (Cleaved-Caspase3), NPC (PAX6) and cell cycle phases (FxCycle) (Table S1). All experiments were performed in triplicates (defined as one hiPSC line cultured in three different wells followed by NPC induction) and included unstained cells and single-color controls (SCCs) using beads or cells. After acquisition and unmixing (Detail in Supplementary File 2, Supplementary File 3, and Analysis Note section), data was pre-processed following field standards to remove low-quality events (Fig. 1, A-D) and enhance signal sensitivity and specificity (Supplementary File 3) (den Braanker et al., 2021; Herzenberg et al., 2006; Park et al., 2020). We used a traditional two-dimensional gating strategy as a benchmark for a more exhaustive, unsupervised assessment and exploratory analysis (Fig. 1 E-L). While downstream analysis used unsupervised algorithms, we defined the minimum expected cluster resolution to inform the parameter selection and workflow. The criteria for minimum expected clusters considered a relatively homogeneous cell population (NPC with varying levels of PAX6 expression) captured at one of three possible stages of the cell cycle (based on markers used, G0/G1, G2, or Mitosis (M)) or undergoing apoptosis (Cleaved-Caspase3). These clusters were expected to have subtle differences in the levels of H3 PTM.

**Figure 1.**
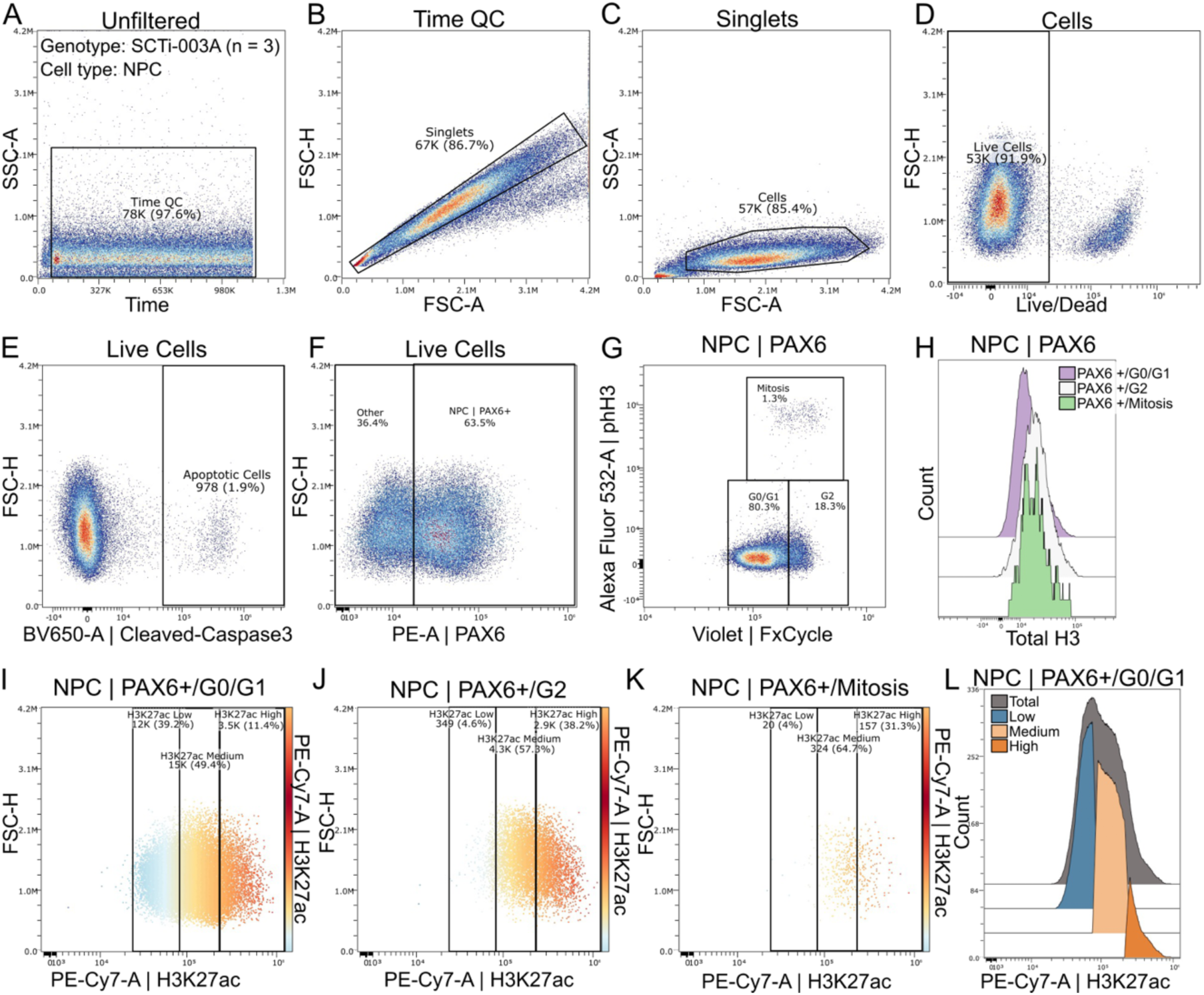
Scatter plots and histograms of manual gating strategies to select high quality events, identify neuronal progenitor cells, their cell cycle status, and histone H3-PTM fluorescence values. (A-D). Cell quality gates to capture single cells recorded at a stable flow rate during acquisition. Subsampling and downstream analyses were performed using Live Cells gate. (E-L) Traditional two-dimensional gating strategy. (E) Within the live cells group, late apoptotic cells (cleaved-Caspase3 positive) are identified. (F) NPCs (PAX6 positive cells) were gated along with cells of low PAX6 expression (“Other”). (G) FxCycle and H3S10ph (pH3-S10) gates were used to identify NPCs in G0/G1, G2, and mitosis. (H) Histogram showing levels of total H3 at G0/G1, G2, and mitosis. Note comparable levels of total H3 during G2 and mitosis. (I-L) Gates to identify low, medium, and high fluorescence intensity values of H3K27ac in each cell cycle phase. Color-scale in panels I-K is used to aid more precise gating of H3 PTM features expected to be distributed as gradients rather than positive-negative cell populations. The same gating strategy was applied to all other H3-PTM assessed. Percent values (%) represent the events of a given gate present in root gate Live cells.

Unlike traditional single-cell transcriptional data analysis, unsupervised clustering of spectral flow data is performed in high-dimensional space, followed by visualization in two-dimensional space. This approach optimizes the high-dimensional data captured by EpiFlow while resolving subtle sub-populations (T et al., 2021). We used the clustering algorithm FlowSOM (Van Gassen et al., 2015) followed by optimized t-stochastic neighbor embedding (opt-SNE) (Belkina et al., 2019). Details about the parameters used in FlowSOM and opt-SNE can be found in Supplementary File 3. We first evaluated the quality of the FlowSOM clusters by comparing them with traditional gates in the opt-SNE two-dimensional space (Fig. 2 A-C). To assess if the obtained clusters were consistent with expected histone patterns (Li et al., 2005; Yiyuan Liu et al., 2017; Ma et al., 2015; Park et al., 2022; Pelham-Webb et al., 2021; Sze et al., 2017; Van Rechem et al., 2021), we inspected the distribution and intensity of each feature by plotting them in a color-continuous scale in the opt-SNE graphs (Fig. 2, D-L). Our analysis resulted in 11 FlowSOM elbow metaclusters that, after careful exploration of all features, were consolidated into 10 clusters (Fig. 2, C; fsom clusters 01-10. For details on merged clusters, see Supplementary File 3). Unsupervised clustering resolved the expected cell populations (Fig.2 D-G) and identified unique cell states or clusters that could potentially be driven by distinct H3 PTM patterns (Fig. 2 C, H-L).

**Figure 2.**
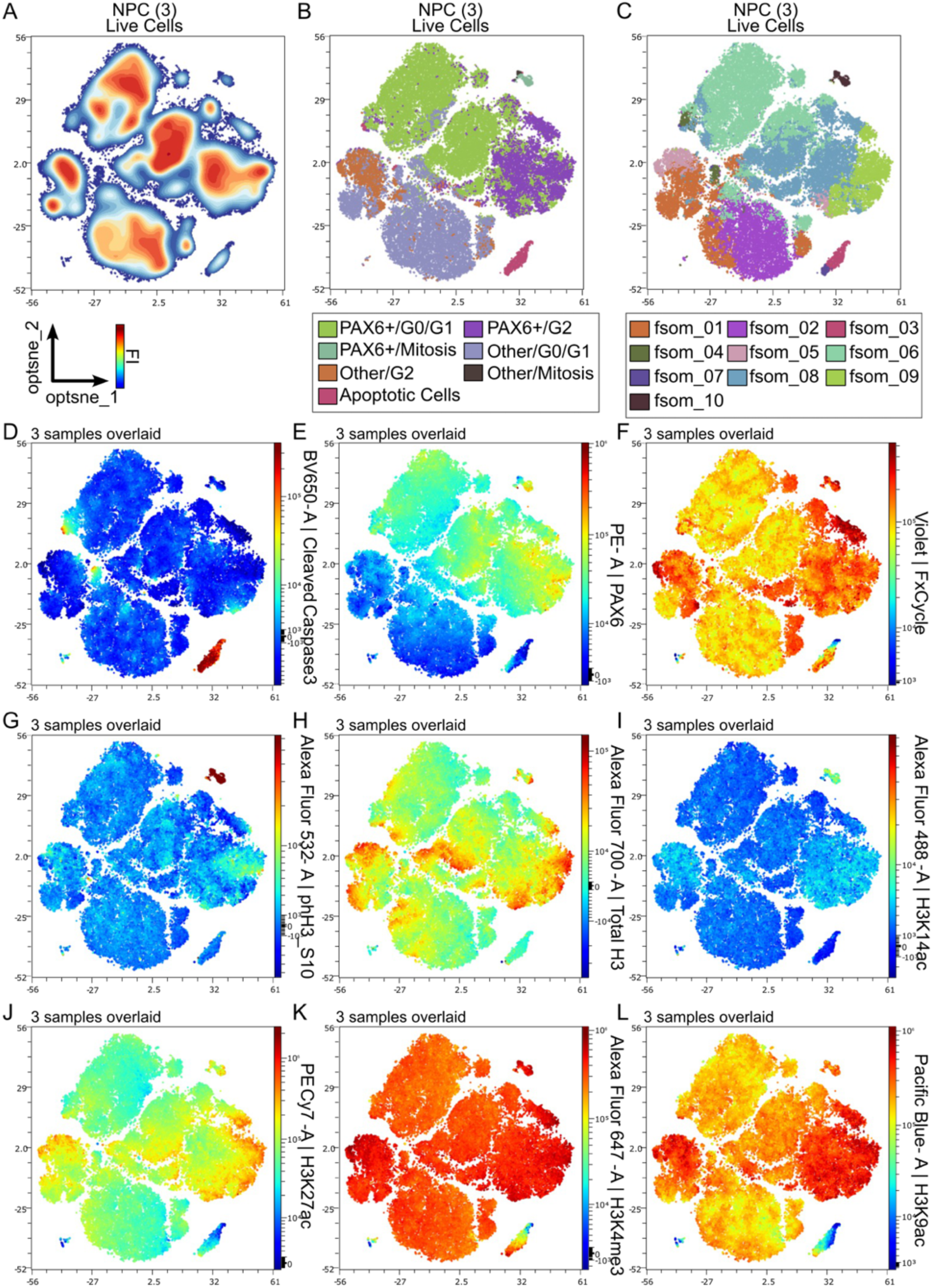
Visualization of manual gating and unsupervised clustering in opt-SNE two-dimensional space. (A) opt-SNE density plot of concatenated NPC samples (n = 3). (B) Cluster identification from traditional two-dimensional gating visualized in the opt-SNE dimensions. (C) Curated FlowSOM clusters visualized in the opt-SNE two-dimensional space. (D-L) Fluorescence intensity values of each feature plotted as a color-continuous scale in opt-SNE twp-dimensional space. The fluorescence intensity (FI) gradient for each feature shows blue for low intensity, green/yellow for medium intensity, and red for high intensity.

Visualization of clusters and features in a two-dimensional space can be useful for initial feature exploration of spectral flow cytometry data. However, the axis and global distances in these plots are meaningless and can be affected by data distortion reported in transcriptional data analysis (Chari and Pachter, 2023; T et al., 2021). Therefore, two-dimensional representations will not accurately reflect the relationships between data points in high-dimensional space. To confirm that each identified cluster represents a distinct and biologically meaningful group of cells, rather than being clustered together by random chance, we analyzed the contribution of H3 PTM patterns in each cluster to determine whether they reliably reflect the cell state or identity. In Figure 3, we evaluated the relative contribution of all features to each cluster, examined the distribution of FI values for all H3 PTMs within each cluster, and analyzed the correlation between features and clusters. In this article, we highlight clusters fsom_08 and fsom_09 as examples. These clusters had similar levels of PAX6 expression (known NPC) and DNA content (FxCycle denoting cells in G2) with subtle differences in FI values for H3 PTMs. Cluster 10 is also discussed due to its unique signature across all features (PAX6 and phH3_S10 double positive cells representing mitotic NPC) (Fig. 3). The analysis of all clusters is available through GitHub Repository associated to this article. Our analysis showed that NPCs in clusters 08 and 09 shared the same five leading features (FxCycle, PAX6, H3K4me3, H3K9ac, and H3K27ac), but PAX6 and H3K27ac contributed differently to each cluster (Fig. 3 A, B). Additionally, while H3K9ac contributed relatively similarly to both clusters (Fig. 3 A, B; third bar), its FI distribution varied between them (Fig. 3 C, E). Our statistical analysis revealed significant differences in the FI and distribution of H3K4me3, H3K9ac, H3K14ac, and H3K27ac across the clusters, suggesting their potential role in distinguishing the biological state of the cells within each cluster (Fig. 3, G-J, Dunn pairwise test for selected clusters. Fig. S5, Krustal-Wallis test for all clusters). When H3 PTMs and cell cycle features are assessed across each cluster, we observed significant differences, suggesting that these clusters represent distinct cellular states of NPCs with varying FI distributions for the analyzed features (Fig. S6).

**Figure 3.**
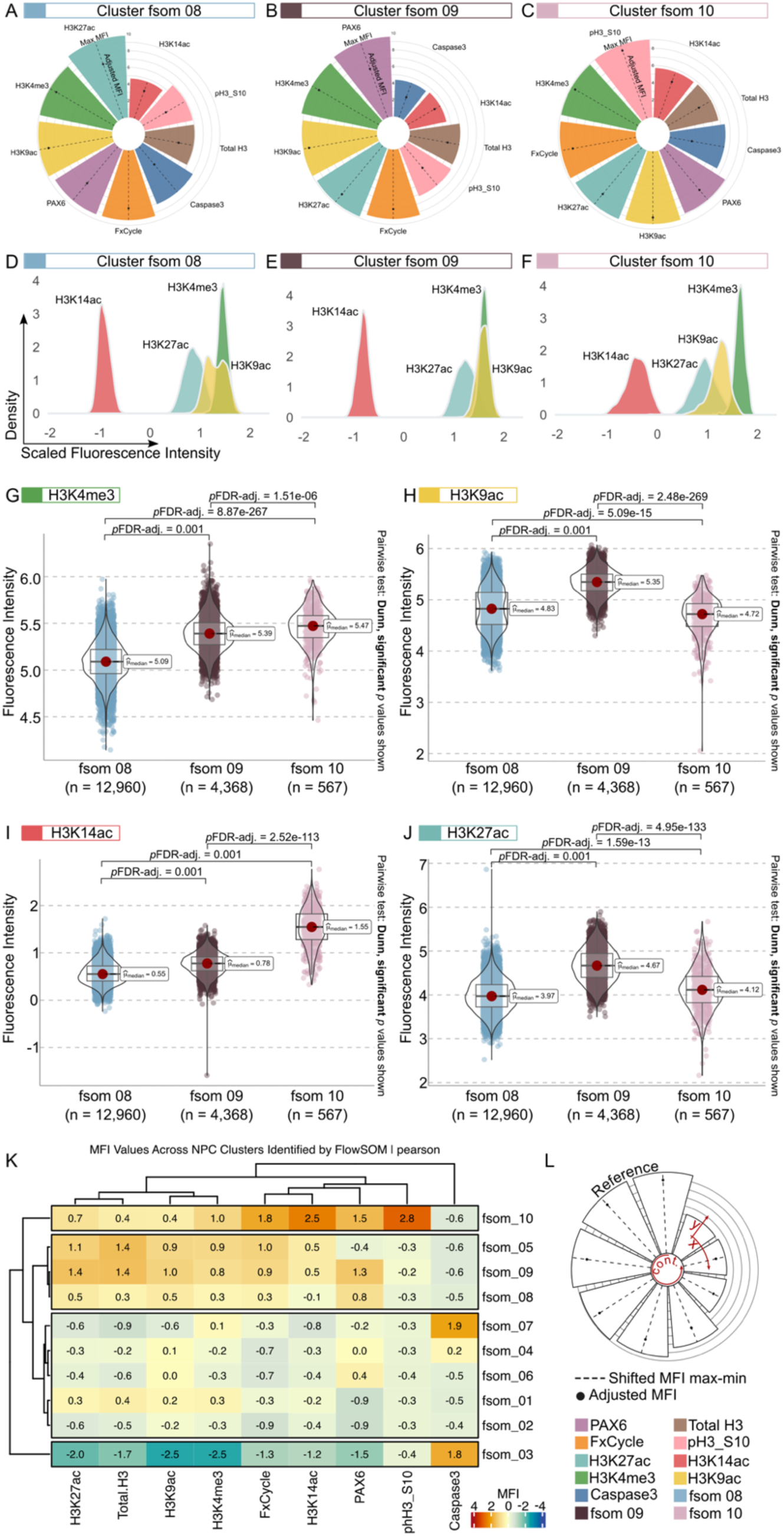
High-dimensional exploration and assessment of H3-PTM profiles across clusters in NPCs. Data analysis and visualization strategies to evaluate feature contributions and distributions to each cluster and assess H3-PTM signatures. (A-C) Circular bar plots displaying the contribution of each feature in selected clusters. Median Fluorescence Intensity (MFI) values were transformed to ensure that all values were positive and greater than zero [MFI shifted = mfi + abs(min(mfi)) + 1]. Values were then summarized by feature (colored bars) within each cluster displaying maximum (outer end of the bar), minimum (inner end of the bar), and median values (shifted MFI, dot in the midline of each bar). Features (bars) are ordered from larger to smaller MFI values in a counterclockwise direction to highlight their contributions to each cluster. (D-F) Density plots showing H3-PTM scaled FI values and distributions in each cluster. (G-J) Violin plots of FI for H3K4me3 (G), H3K9ac (H), H3K14ac (I), and H3K27ac (J) across fsom_08, fsom_09, and fsom_10. Dunn pairwise test, statistically significant *p*-values shown. (K) Heatmap displaying the MFI values across 10 NPC clusters. To assess the correlation between H3-PTM, cell cycle status, and expression levels of PAX6, clustering of scaled MFI values (displayed within the heatmap) was performed using the Pearson correlation distance method. (L) Reference cartoon to support the interpretation of circular bar plots and color-code of each feature.

## Discussion

In this study, we developed and validated EpiFlow, a spectral flow cytometry protocol that profiles histone PTMs and cell state markers in whole cells. This analysis provides unprecedented insights into the multidimensional complexity of cellular state. EpiFlow overcomes key limitations of traditional flow cytometry by enabling high-resolution analysis of epigenetic modifications while preserving cellular context. This approach resolved a gradient of PAX6-expressing NPC clusters (Fig. 2, 3), correlating their varying fluorescence intensities of histone PTMs with unique features like specific cell cycling phases, a level of resolution currently unattainable by conventional methods. Additionally, we visualized feature contributions (Fig. 3 A-C) and distributions (Fig. 3 D-F) to identify the individual and collective impacts of each fluorophore to the overall cell state. These results suggest that specific molecular markers (e.g., histone PTMs, cell cycle, or lineage transcription factors) independently and interactively shape cellular state (Tarazona and Pourquié, 2020; Torcal Garcia and Graf, 2021; Wang and Higgins, 2013). By plotting fluorescent readout distributions and analyzing histogram shapes, we revealed feature variability and heterogeneity within cell populations. Differences in histogram structures can identify subpopulations with distinct molecular characteristics, subtle shifts in expression levels, and potentially rare states within broader populations. This analysis also presented correlations and plausible dependencies between markers (Fig. 3 K), permitting deeper inquiries into how specific features might dynamically regulate cell states. This approach enhances our ability to characterize cellular heterogeneity and provides a framework to understand the interplay between molecular events and global cellular outcomes, advancing both the basic biology and translational research of epigenetic regulation.

In this work, we demonstrate that EpiFlow is adaptable across multiple flow cytometry platforms and cell lines. Our findings suggest that subtle variations in histone PTMs across the cell cycle contribute to within-population heterogeneity, even in seemingly homogeneous cell cultures, such as hiPSC-derived NPCs (Figs. 1, 2). While variability in differentiation or induction efficiency is a well-documented challenge in stem cell research (Cuomo et al., 2020; Nishizawa et al., 2016; Zakrzewski et al., 2019), our results suggest that heterogeneity in histone PTM deposition and cell cycle dynamics may also underlie variability in differentiation outcomes. Importantly, our drug validation studies and use of a functionally deficient methyltransferase hiPSC line (KMT2D10.4) confirm that this variability is not attributable to differences in instrumental sensitivity (Fig. S2, S3, S4). Future experiments could explore the use of modulators targeting these features to enhance differentiation efficiency and reproducibility (Ciceri et al., 2022; Yang et al., 2016). Additionally, while both manual gating and unsupervised clustering of spectral flow data effectively identify similar broad cell populations, unsupervised clustering using the FlowSOM algorithm uncovered unique cell states characterized by more complex, multidimensional features. However, careful consideration of the underlying biology is essential, as the algorithm can generate additional clusters based on artifacts such as debris. For example, former Cluster 03, initially identified due to an outstanding debris feature contribution, was later merged with the apoptotic cluster upon closer examination (File S3). Future work should expand the application of EpiFlow to other cell types and experimental systems to further validate its utility.

Our spectral flow cytometry data reveal distinct chromatin modification profiles across NPC Clusters 08, 09, and 10 (Fig. 3), suggesting these clusters could represent different stages or functional states within the NPC population. Cluster 10 is characterized by increased cycling marks (pH3 S10 and FxCycle stain) along with a strong H3K14ac signal and a broad histogram spread, potentially indicative of active acetylation dynamics linked to open chromatin states required for transcription during the cell cycle (Kishkevich et al., 2019; Wang et al., 2012) (Fig. 3 C, F, G-J). The weaker and more variable H3K9ac and H3K27ac signals suggest transient or heterogeneous genomic regulatory activity, while the strong H3K4me3 signal could reflect the maintenance of transcriptionally active marks (Beacon et al., 2021; Benayoun et al., 2014). Cluster 9 shows overlapping strong signals of H3K4me3 and H3K9ac, suggesting a maturing state, possibly as these cells prepare for differentiation or a shift in transcriptional programs (Kim et al., 2011) (Fig. 3 B, E, G-J). Indeed, H3K9ac levels is known to rise in maturing NPCs (Du et al., 2017; Qiao et al., 2015). The upward shift in the H3K27ac histogram spread compared to Cluster 10 suggests increased enhancer activity for processes that could be related to lineage-specific gene regulation (Rubin et al., 2017). The minimal H3K14ac signal could reflect reduced global chromatin opening, signaling the initiation of lineage commitment or the transition out of the proliferative state (Baell et al., 2018; Yang et al., 2022). Notably, Cluster 09 also exhibits stronger PAX6 expression, which is also observed in maturing NPCs (Sansom et al., 2009), further supporting its description as a potentially maturing progenitor cell state. Interestingly, Cluster 08 again displays a strong H3K4me3 signal along with two distinct peaks in the H3K9ac histogram, indicating potential heterogeneity within this feature (Fig. 3 A, D, G-J). One H3K9ac peak retains a histogram profile similar to Cluster 10’s H3K9ac peak, while the other peak aligns more closely with histogram profile of the maturing Cluster 09. This duality suggests that Cluster 08 could represent NPCs transitioning between cycling and maturing states. The weak and broad H3K27ac signal may reflect less uniform enhancer activation, while minimal H3K14ac levels again point to potentially reduced chromatin acetylation as cells stabilize for lineage-specific gene expression.

These findings support the notion that histone PTMs are dynamic across the cell cycle and unique to cell state. EpiFlow has demonstrated its utility in characterizing such epigenetic profiles, which, in line with previous research showing that histone PTMs predict cell fate acquisition (Karlić et al., 2010; Trevino et al., 2021), could also indicate future cell potential if these NPCs were to be differentiated in distinct neural maturation media. However, the consistently weaker H3K27ac signal raises the possibility of suboptimal antibody staining, which will require troubleshooting in future experiments. While EpiFlow can detect subtle differences within fairly homogenous cell populations, assigning labels or identities to clusters is limited by the markers used in the panel, such as PAX6 or DNA content for cell cycle phases. Furthermore, EpiFlow does not provide genomic information about the histone marks nor link these modifications to specific chromatin regions or regulatory elements. Therefore, inferring the biological relevance of a given cluster based on its H3-PTM profile should be regarded cautiously without downstream analyses and validation. Cross-validation using ChIP-seq can confirm the genomic localization and functional relevance of these histone modifications, while temporal analyses of NPC differentiation can validate the progression of chromatin states and cell identities over time, reinforcing the biological transitions inferred from this analysis.

EpiFlow offers a significant contribution to epigenetic research by bridging the gap between traditional cytometry and high-dimensional omics approaches. While current clinical applications often rely on DNA methylation as a biomarker for disease diagnosis and prognosis (García-Giménez et al., 2017; Hao et al., 2017), EpiFlow introduces histone PTMs as a complementary flow cytometry tool that provides additional layers of information about chromatin dynamics. Additionally, integrating spectral flow cytometry with other omics approaches (Zhou et al., 2024) could further enhance insights into epigenetic regulation and its role in development and disease. The method described here can be adapted to study diverse cell types, samples, and biological systems, including those in cancer biology, organoid systems, biopsies, and immunological studies, broadening its relevance and utility. Moreover, through the associated GitHub repository, we provide a reproducible and accessible workflow for the exploratory and statistical analysis presented in this article. In conclusion, EpiFlow provides a powerful platform for advancing our understanding of chromatin biology and has the potential to transform both basic research and clinical applications in epigenetics.

## Materials and methods

The SCTi003-A (RRID:CVCL_C1W7) and WTC11 (UCSFi001-A, RRID:CVCL_Y803) control hiPSC lines were used throughout this study. KMT2D loss-of-function (LOF) hiPSC line was generated through CRISPR/Cas9 editing of exon 2 in the *KMT2D* gene. This resulted in a homozygous 2 bp deletion in the wildtype WTC11 iPSC line, creating the isogenic KMT2D-LOF model (KMT2D10.4). Room temperature refers to 22 °C. All hiPSC lines were karyotyped and confirmed to be free of mycoplasma. This protocol can be applied to samples run on traditional flow cytometers, including the Stratedigm S1000EXi benchtop flow cytometer. For this, the panel design process should be repeated to ensure that the fluorochromes selected are compatible with the instrument. Details on the Stratedigm assays are in Table S2. All experiments were conducted as n=3 biological triplicates. ChatGPT4o (AI) was used to assist in text refinement and code optimization.

### Cell culture

hiPSCs were grown on 6-well plates (Corning) coated with 2D Matrigel (Corning) in mTeSR Plus media (Stem Cell Technologies) supplemented with Primocin (InvivoGen). Once the cells were ∼70% confluent, the media was replaced with STEMdiff SMADi Neural Induction media (Stem Cell Technologies) supplemented with Primocin. The neural induction was performed following the manufacturer’s protocol. Cells were subcultured at 70% confluency to maintain continuous mitosis events by day 7. All cultures were incubated at 37°C in a 5% CO2 incubator with daily media changes.

### Drug Treatments

NPCs were cultured to 70% confluency and treated with three different concentrations of Trichostatin A (TSA; 0.5μM, 1μM, 10μM) or A485 (1μM, 10μM, 20μM). SCTi-003A, WTC11 and KMT2D10.4 hiPSCs were also grown to 70% confluency and treated with TSA (0.5μM, 1μM, 2μM), Vorinostat (SAHA; 0.5μM, 0.75μM,1μM), OICR-9429 (10μM, 20μM, 50μM) or CPI-455 HCl (10μM, 20μM, 50μM). Cells were treated for 4 hours with either the drug or DMSO, then harvested and processed according to the flow cytometry protocols outlined above.

### Sample preparation

On day 7 NPCs were gently detached and resuspended in a round bottom 96-well plate. Cell pellets were stained with fixable viability dye (BD) for 15 minutes on a rotating platform at room temperature. Cells were then fixed in 4% PFA (Thermo Fisher) for 20 minutes at room temperature. For permeabilization, cells were incubated with 100% ice-cold methanol (Fisher Chemical) for 15 minutes in a -20°C freezer. For blocking, cells were incubated with 5% BSA for 30 minutes on a rotating platform at room temperature. Cells were then incubated with the appropriate antibody cocktail for 1 hour on a rotating platform at room temperature. Finally, cells were stained with FxCycle Flow Ready (Invitrogen) for 30 minutes on a rotating platform at room temperature. FxCycle was not removed from the solution prior to running on the cytometer. All incubations were performed with a shaking speed of ∼80 rpm. Between each step (except FxCycle), cells were washed once in FACs buffer (0.5% BSA, Sigma-Aldrich). All centrifugal spins were performed at 4°C for 5 minutes. Following the spin, a quick and swift decanting movement was used to ensure that sample loss and cross-contamination were prevented. Single color controls (SCCs) were prepared using beads (Invitrogen) or cells. Cells were used for all dye-based staining. After dyes and antibodies were added, the samples were protected from light using foil.

### Spectral flow cytometry assay

Data was acquired on a 5L Cytek Aurora analyzer (Cytek Biosciences). Before acquiring the data, ensure that each sample is well-mixed by pipetting immediately before running the assay. Ensure that the Aurora completed the warmup/start-up initialization and passed the quality control check. Run the spectral flow cytometry assay according to user-defined settings. We utilized the following acquisition settings for NPCs: Events to record: 10,000,000, Stopping time (sec): 10,000, Stopping volume (μL): 150, Flow rate: High, FSC: 40, SSC: 20.

### Data unmixing

Acquired data was unmixed with SpectroFlo v2 (Cytek Biosciences) without autofluorescence removal. Unstained cells and SCCs (beads or cells) were used as references for unmixing, as detailed in Table S1. After unmixing, both SCCs and fully stained samples were manually inspected for potential unmixing artifacts in NxN plots (File S2) following recommended guidelines for spectral flow cytometry panel optimization (Ferrer-Font et al., 2021).

### Gating, clustering and dimensionality reduction

The data analysis workflow for gating and dimensionality reduction was built using the cloud-based OMIQ analysis platform (File S3). To ensure all events were properly scaled, each feature was transformed using arcsinh (File S3). Three gating strategies were implemented to optimize signal separation, exclude artifacts or debris, and identify distinct cell populations, ensuring accurate downstream analysis.

*Event rate and cell quality gating:* First, time gating was applied to select events recorded with a stable flow rate. This was followed by additional gating to isolate singlets, exclude debris, and discriminate live and dead cells. Subsequent gating and analyses were performed within the live cell population.

*Manual gating as a benchmark:* This strategy involved the identification of apoptotic cells (cleaved Caspase-3 positive cells), neuronal progenitor cells (PAX6 positive cells), and determining cell cycle status (G0/G1-G2: FxCycle; mitosis: pH3-S10 positive cells). To account for differences in fluorescence intensity signal observed in H3K9ac, H3K4me3, H3K14ac, and H3K27ac features, distinct gates were applied targeting events with low, medium, and high fluorescence intensity within each cell cycle phase. Changes in global H3 were assessed across G0/G1, G2, and mitosis. Fluctuations in H3-PTM fluorescence intensity, potentially driven by H3 replication-dependent biogenesis (Armstrong and Spencer, 2021), were evaluated by applying specific gates for each histone mark within the Total H3 gate (Fig. S7). After manual gating, the subset of fully stained files was retained and the maximum number of live cells was randomly subsampled from each replicate.

*Unsupervised clustering and gating:* Unsupervised clustering was performed using FlowSOM (Van Gassen et al., 2015). The resulting elbow meta-clusters (“fsom clusters”), representing distinct groupings of data that share similar characteristics, were embedded as gates for further analysis and comparison with manual gating. Dimensionality reduction was performed using optimized T-distributed stochastic neighbor embedding (opt-SNE) (Belkina et al., 2019). Data was exported from OMIQ and is available in the GitHub repository associated with this article. Please refer to File S3 for a detailed workflow, including feature selection, analysis parameters, and random seed details for each algorithm.

### Data exploration and visualization

The data exploration and visualization workflow was built in R using the renv package (Renv v1.0.7), which creates reproducible R environments and ensures data analysis reproducibility. This workflow provides a graphical representation of the main features contributing to each FlowSOM cluster. Methods and statistical analyses are described in the figure legends. A comprehensive analysis and data visualization code, including input data, data processing steps, and visualization rationale, is available at [https://github.com/SerranoLab/Golden_etal_2024].

### Reagents

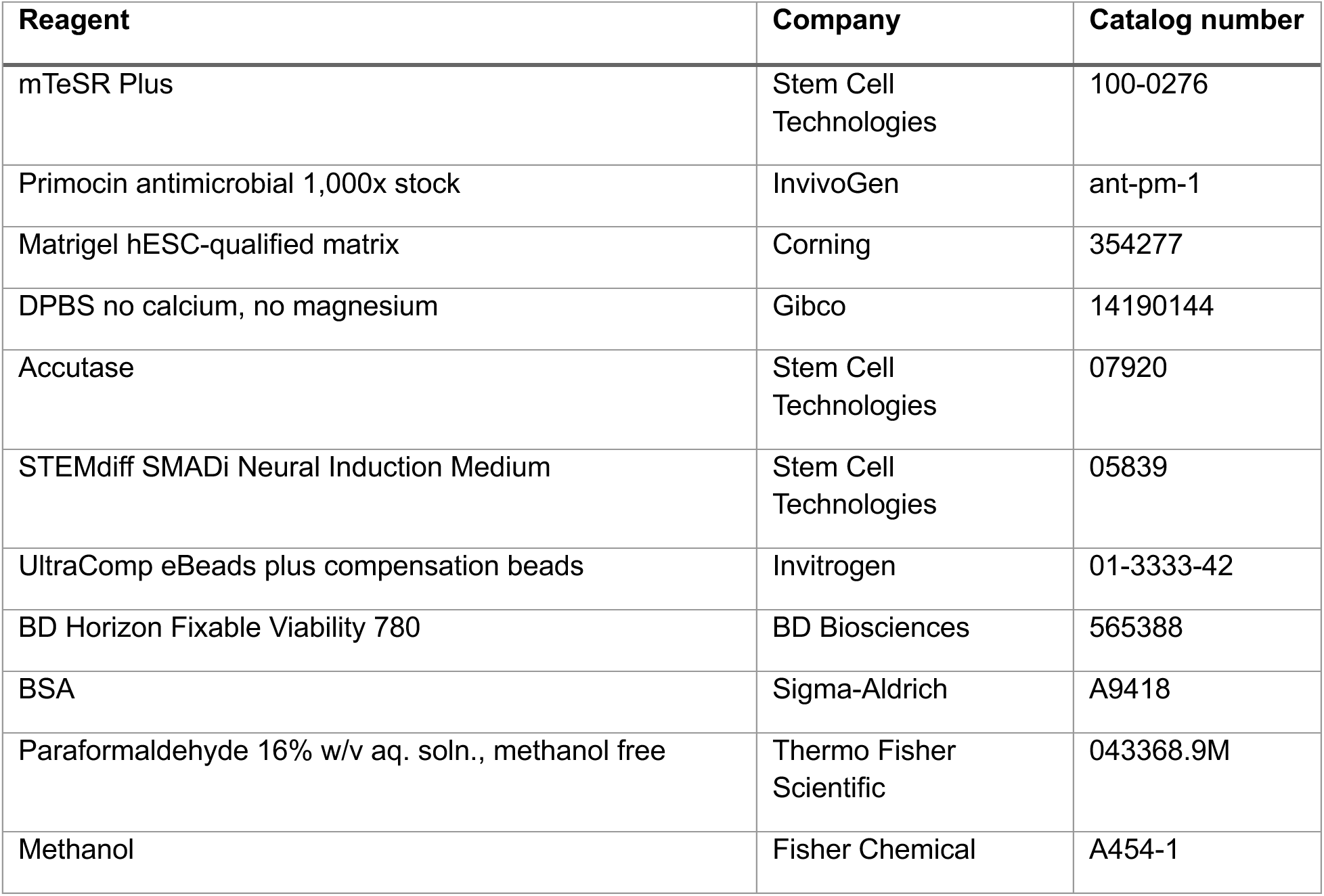
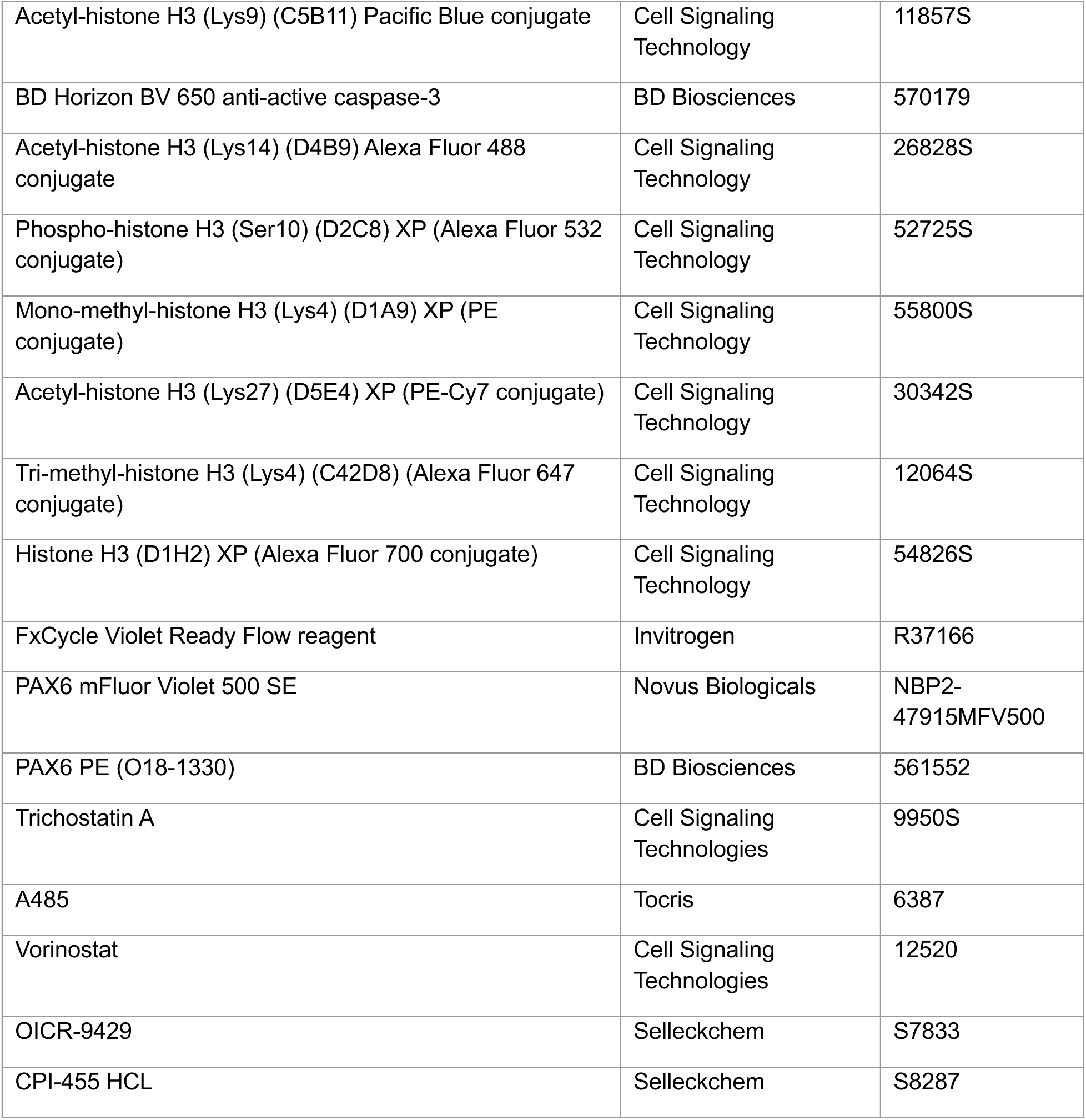

## Acknowledgments

The authors would like to thank Dr. Shari Brezinsky from Boston University Flow Cytometry Core Facility for instrumental support, Irena Feng for supporting initial method optimization, and Sandra Sulser for sample processing support.

This research was supported by Startup Funds to MAS and the Boston University Clinical & Translational Science Institute (CTSI) TL1 Pre-Doctoral Fellowship in Regenerative Medicine (TL1TR001410) to CSG. WTC11 and KMT2D-/- cell lines were provided by grant UM1 HL098160 to HJY from the National Heart, Lung, and Blood Institute. The contents of this work are solely the responsibility of the authors and do not necessarily represent the official views of the NIH.

## Data and resource availability

Data analysis code and details of R packages used are available at [https://github.com/SerranoLab/Golden_etal_2024]. Raw flow cytometry and unmixed spectral flow cytometry data will be made available in FlowRepository.

---

**Figure S1:**
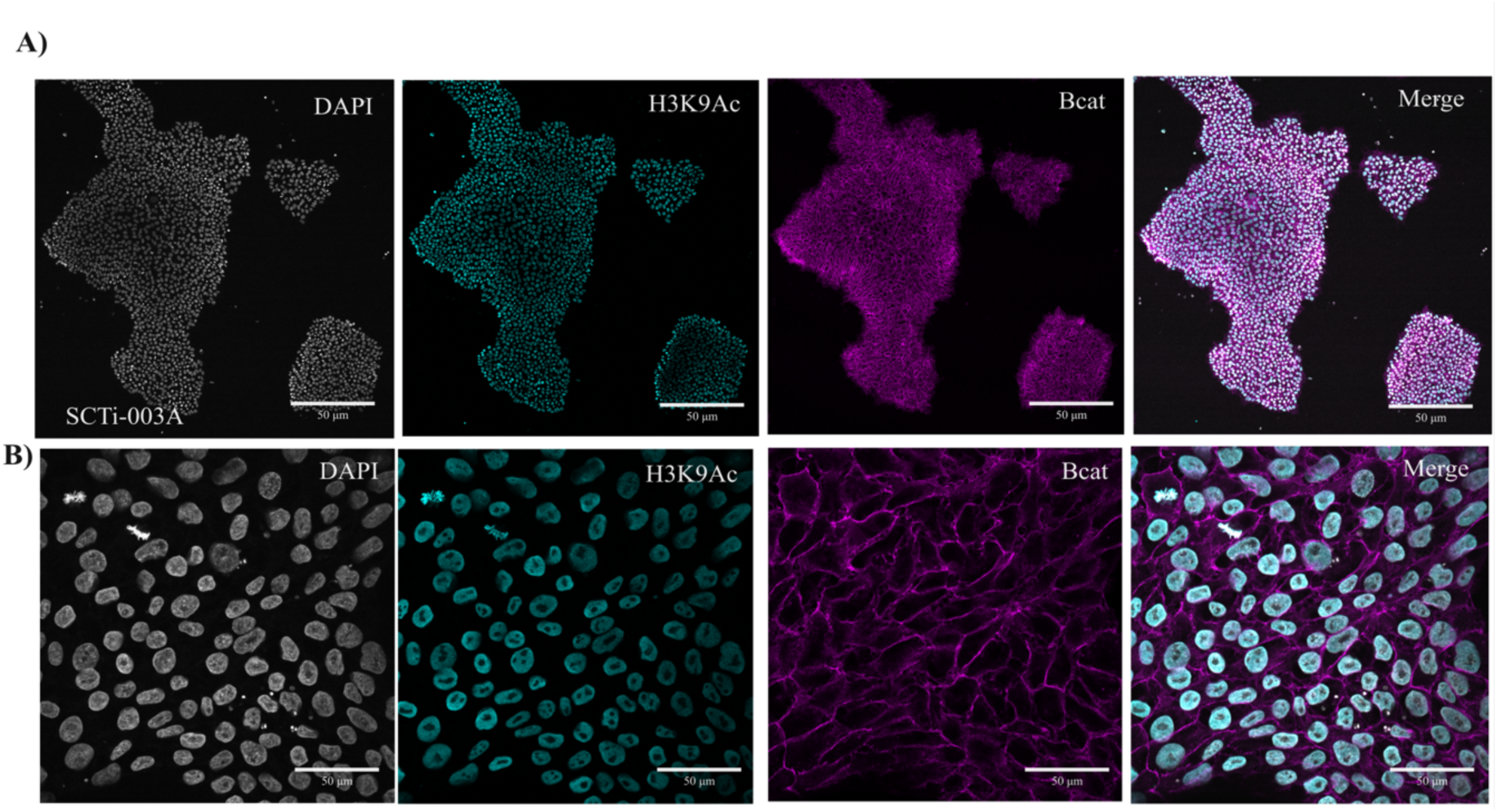
Validation of nuclear and membrane compartment staining in SCTi-003A hiPSCs. **Immunofluorescent staining validation of cell membrane integrity and nuclear antigen localization.** Confocal microscopy images of SCTi-003A hiPSCs. Cell colonies were fixed on glass coverslips and stained for β-catenin (Bcat) and histone 3 lysine 9 acetylation (H3K9ac) to assess membrane integrity and nuclear antigen location, respectively. (A) 10x magnification and (B) 63x magnification confocal images of SCTi-003A hiPSC colonies. Images were acquired on a Zeiss LSM 710 confocal microscope. Scale Bar: 50 μM.

**Figure S2.**
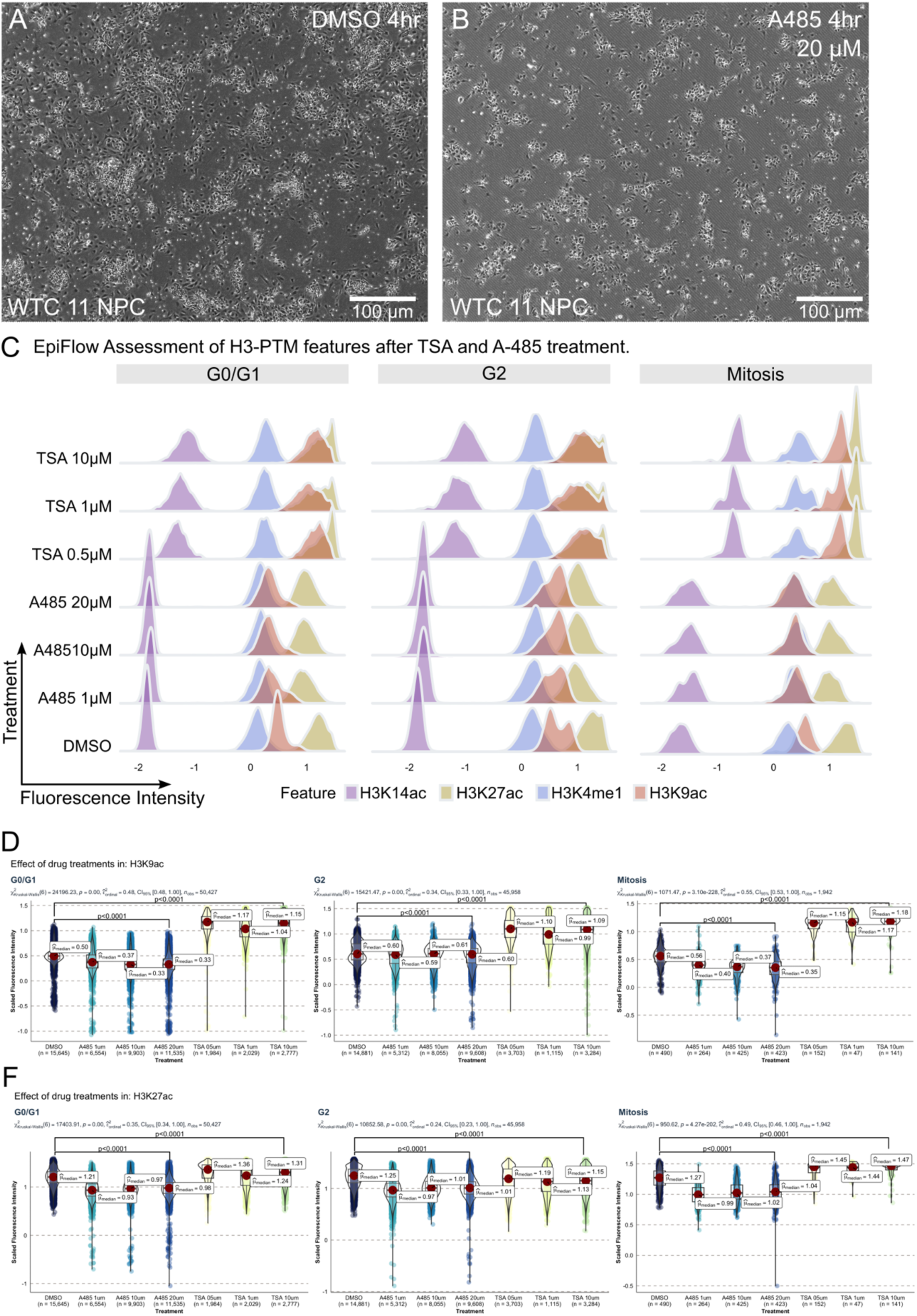
Drug validation assay of hiPSC-derived NPCs using spectral flow cytometry protocol. **Validation of spectral flow cytometry protocol using selective epigenetic protein inhibitors.** Spectral flow cytometry analysis of WTC11 NPC treated with TSA and A485. TSA is a histone deacetylase I/II inhibitor, resulting in increased histone acetylation. A485 is a histone acetyltransferase inhibitor that targets p300/CBP, resulting in decreased histone acetylation. For each drug treatment, cells were treated for 4 hours with DMSO (A), A485 (B), or TSA. A representative brightfield image from one concentration of A485 is shown. Superimposed density histogram plots for H3K9ac, H3K27ac, H3K4me1, and H3K14ac features across all drug concentrations display how the signal is distributed within each group throughout the cell cycle (C). Violin plots of the same groups confirm that the drugs were able to significantly alter H3-PTM acetylation levels (D, E). Replicates n=3 per group. Scale bar: 100 um. TSA: Trichostatin A. NPC: neural progenitor cells.

**Figure S3.**
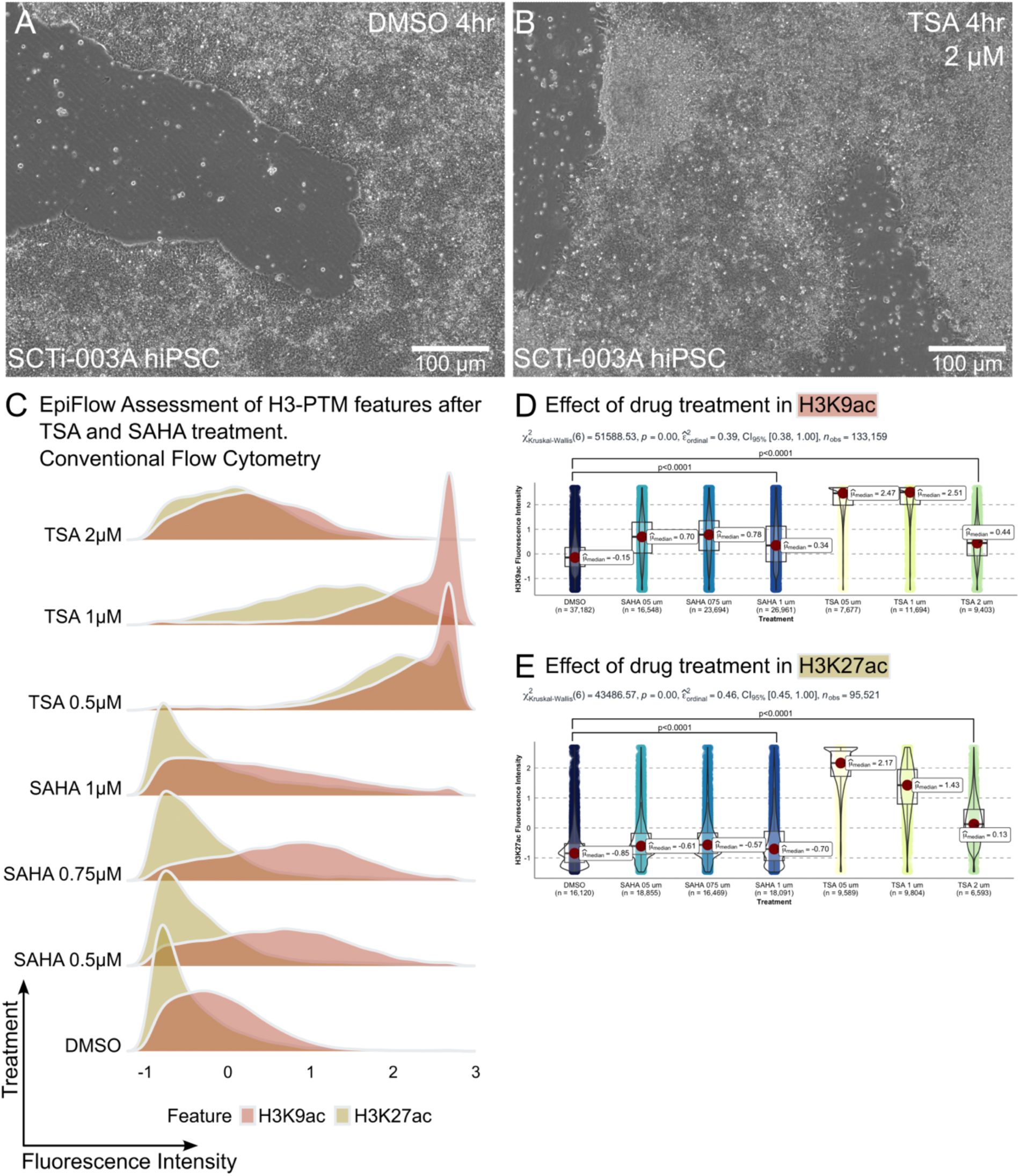
Drug validation assay of hiPSCs using conventional flow cytometry protocol. **Validation of conventional flow cytometry protocol using epigenetic protein inhibitors.** Conventional flow cytometry analysis of SCTi-003A hiPSCs treated with TSA and Vorinostat (SAHA). TSA and SAHA are histone deacetylase I/II inhibitors, resulting in increased histone acetylation. For each drug treatment, cells were treated for 4 hours with DMSO (A), TSA (B), or SAHA. A representative brightfield image from one concentration of SAHA is shown. Superimposed density histogram plots for H3K9ac and H3K27ac features across all drug concentrations show how the signal is distributed in each group (C). Violin plots of the same groups confirm that the drugs were able to significantly alter H3-PTM acetylation levels (D, E). Replicates n=3 per group. Scale bar: 100 um. TSA: Trichostatin A. SAHA: Vorinostat. hiPSC: human induced pluripotent stem cells.

**Figure S4.**
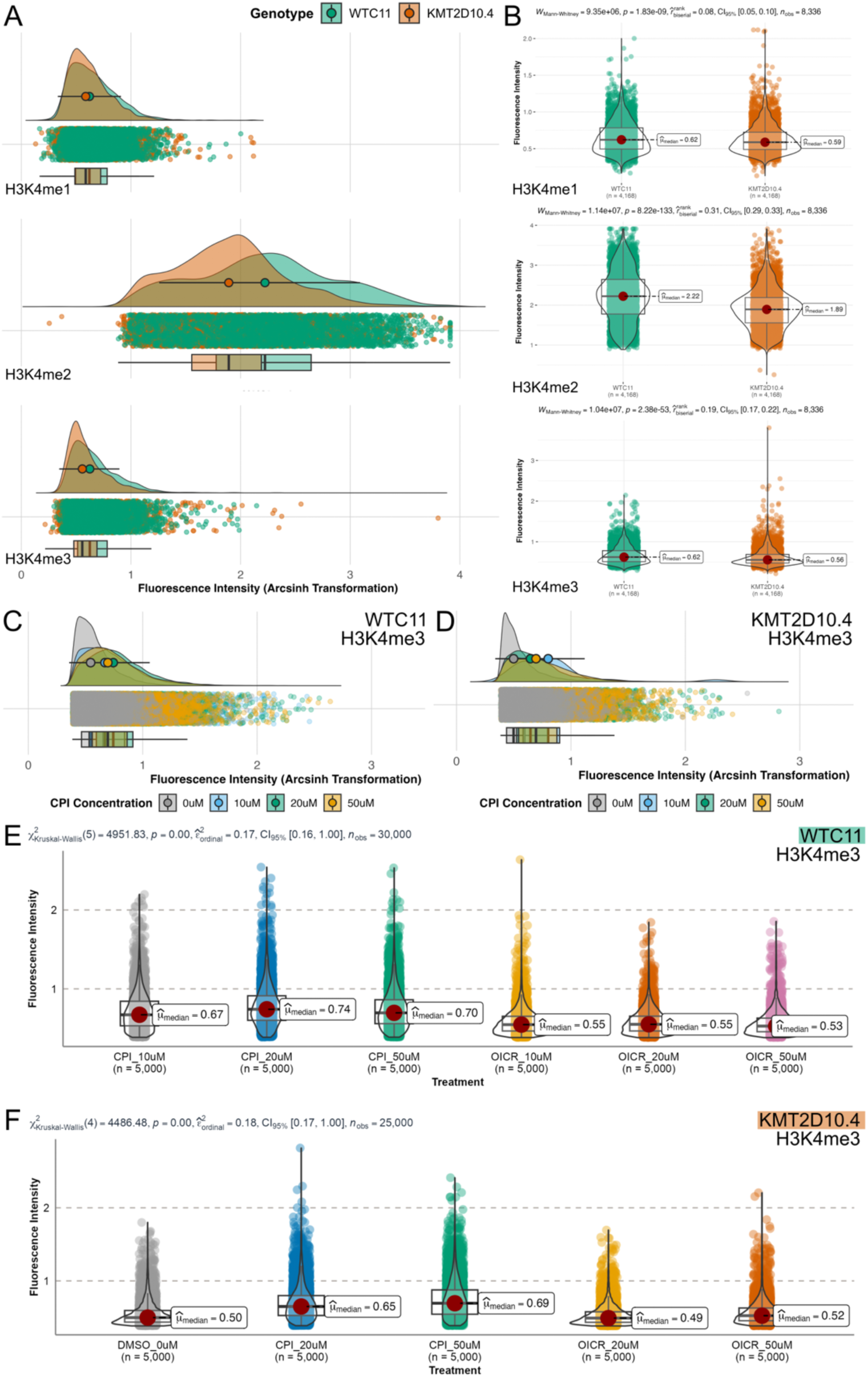
Drug validation assay targeting H3K4 methylation in hiPSCs using conventional flow cytometry. EpiFlow Detects Subtle Changes in H3K4 Methylation Marks. EpiFlow sensitivity was validated using a KMT2D loss-of-function (LOF) hiPSC line, which was generated through CRISPR/Cas9 editing of exon 2 in the KMT2D gene. This resulted in a homozygous 2 bp deletion in the wildtype WTC11 iPSC line, creating the KMT2D-LOF model (KMT2D10.4). WTC11 and KMT2D10.4 hiPSCs were incubated with DMSO or varying concentrations of OICR (H3K4me antagonist), or CPI (H3K4me agonist), for six days in culture. Data was acquired using conventional flow cytometry, preprocessed in OMIQ (scaling, gating, subsetting, subsampling), and analyzed in R. All plotted events correspond to H3K4 methylation within the H3-positive gate. (A-B) Rain plots (A) and violin plots (B) comparing fluorescence intensities of H3K4me1, H3K4me2, and H3K4me3 in KMT2D10.4 mutants versus WTC11. Statistically significant reductions in signal intensity and event counts are observed in the mutant line (Mann-Whitney U test, *p < 0.05* for all marks). (C-D) H3K4me3 levels in WTC11 (C) and KMT2D10.4 (D) hiPSCs treated with CPI (10 μM, 20 μM, and 50 μM) and OICR (10 μM, 20 μM, and 50 μM). (E-F): Violin plots and statistical analysis of H3K4me3 levels in WTC11 (E) and KMT2D10.4 (F) after treatment with CPI and OICR (Kruskal-Wallis test, p < 0.05; 3 biological replicates).

**Figure S5.**
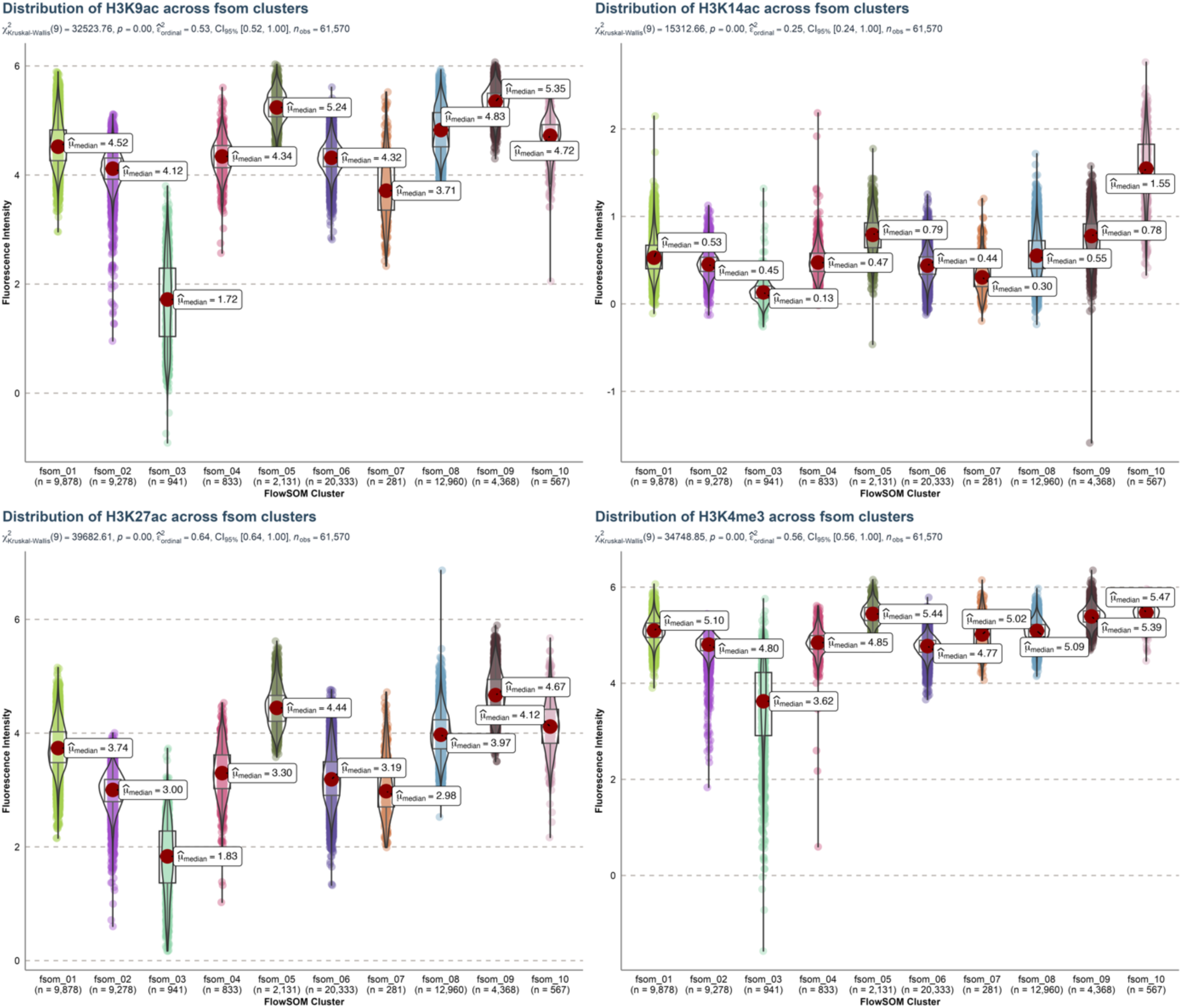
Distribution of H3 PTM fluorescence intensity in NPCs across 10 FlowSOM clusters. Violin plots of fluorescence intensity distributions for H3K9ac, H3K14ac, H3K27ac, and H3K4me3 across 10 clusters. Median values for each cluster are indicated with red dots, and their corresponding numerical values are labeled. Krustal-Wallis test shows significant differences in each feature assessed H3K9ac χ^2^(9) = 32523.76, *p*<0.0001; H3K14ac χ^2^(9) = 15312.66, *p*<0.0001; H3K27ac χ^2^(9) = 39682.61, *p*<0.0001; H3K4me3 χ^2^(9) = 34748.85, *p*<0.0001.

**Figure S6.**
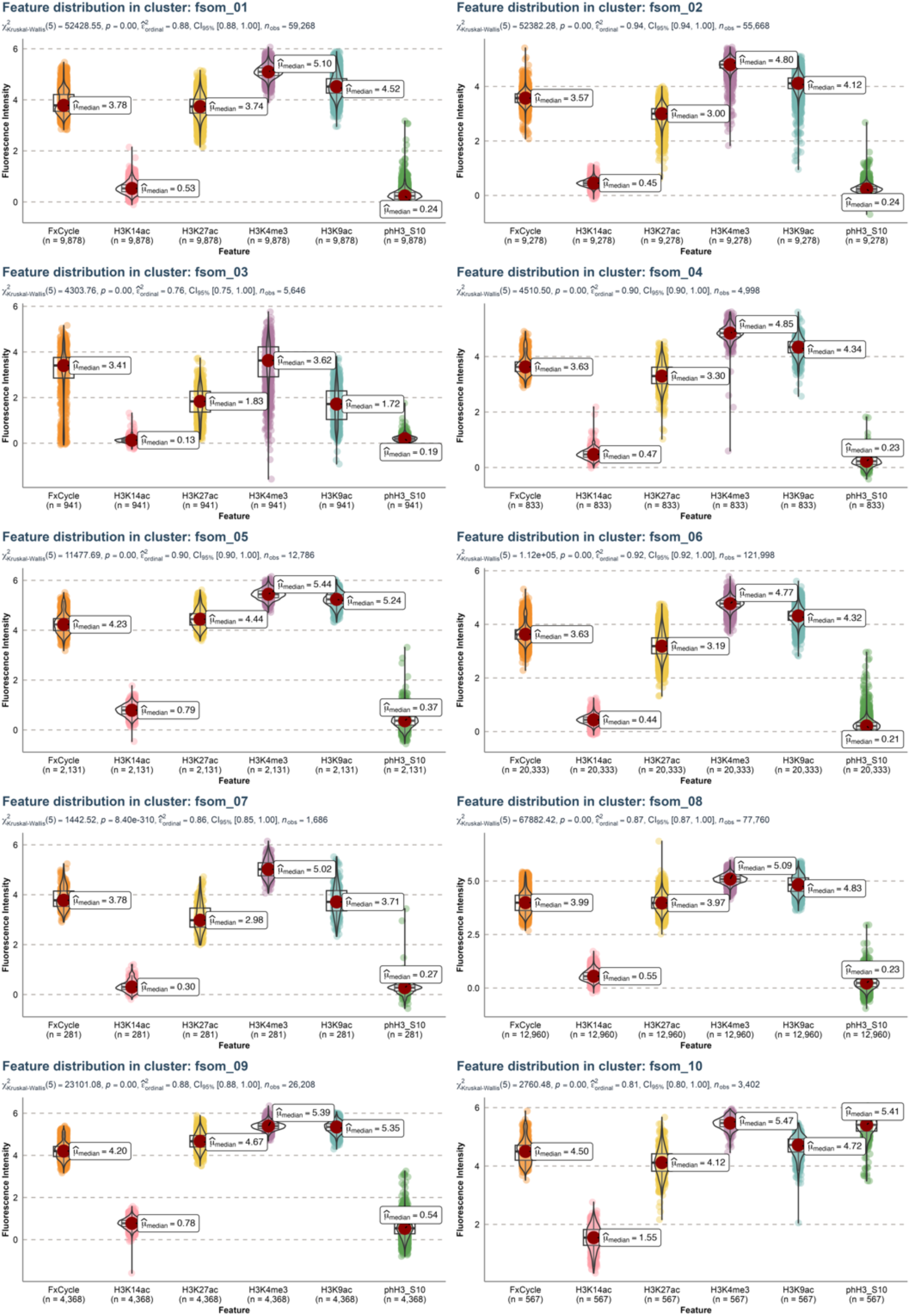
Assessment of H3 PTM signatures and cell cycle markers within clusters. Violin plots representing fluorescence intensity distributions of FxCycle, H3K14ac, H3K27ac, H3K4me3, H3K9ac, and phH3_S10 across 10 clusters (fsom_01 to fsom_10). Boxplots within each violin plot show the interquartile range and median fluorescence intensity values (red spot). Kruskal-Wallis test was applied to compare fluorescence intensities of all features within each cluster. Dunn’s pairwise test was performed for post-hoc analysis. Sample sizes (n) are indicated for each feature in each cluster.

**Figure S7.**
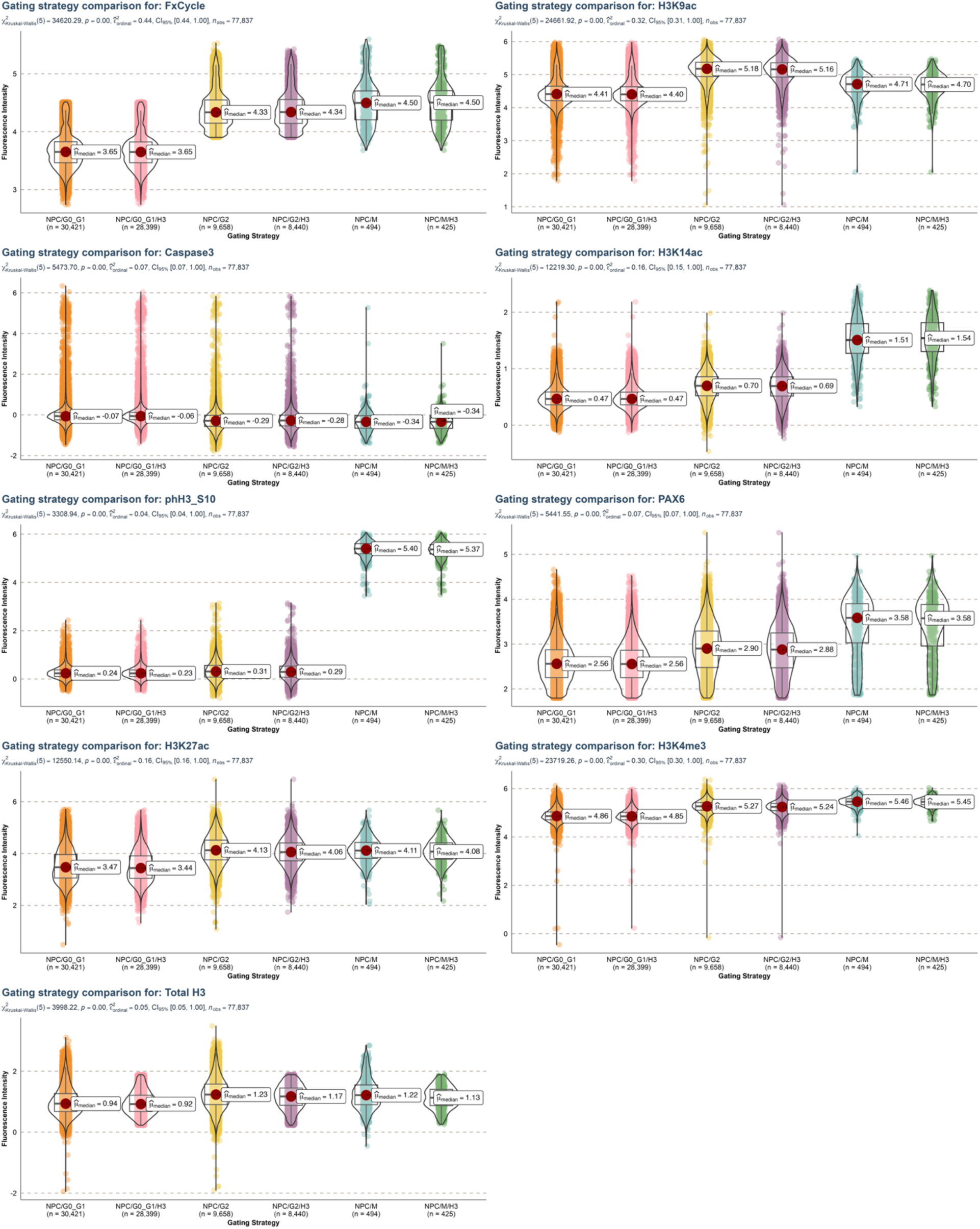
Comparison of Gating Strategies for H3 PTMs and Cell Cycle Markers Across G0/G1, G2, and Mitosis in NPCs. Violin plots of all features across gated cell populations in different cell cycle phases. Kruskal-Wallis test statistics and p-values are provided for each feature. Dunn pairwise comparisons are available in GitHub Repository.

**Figure S8.**
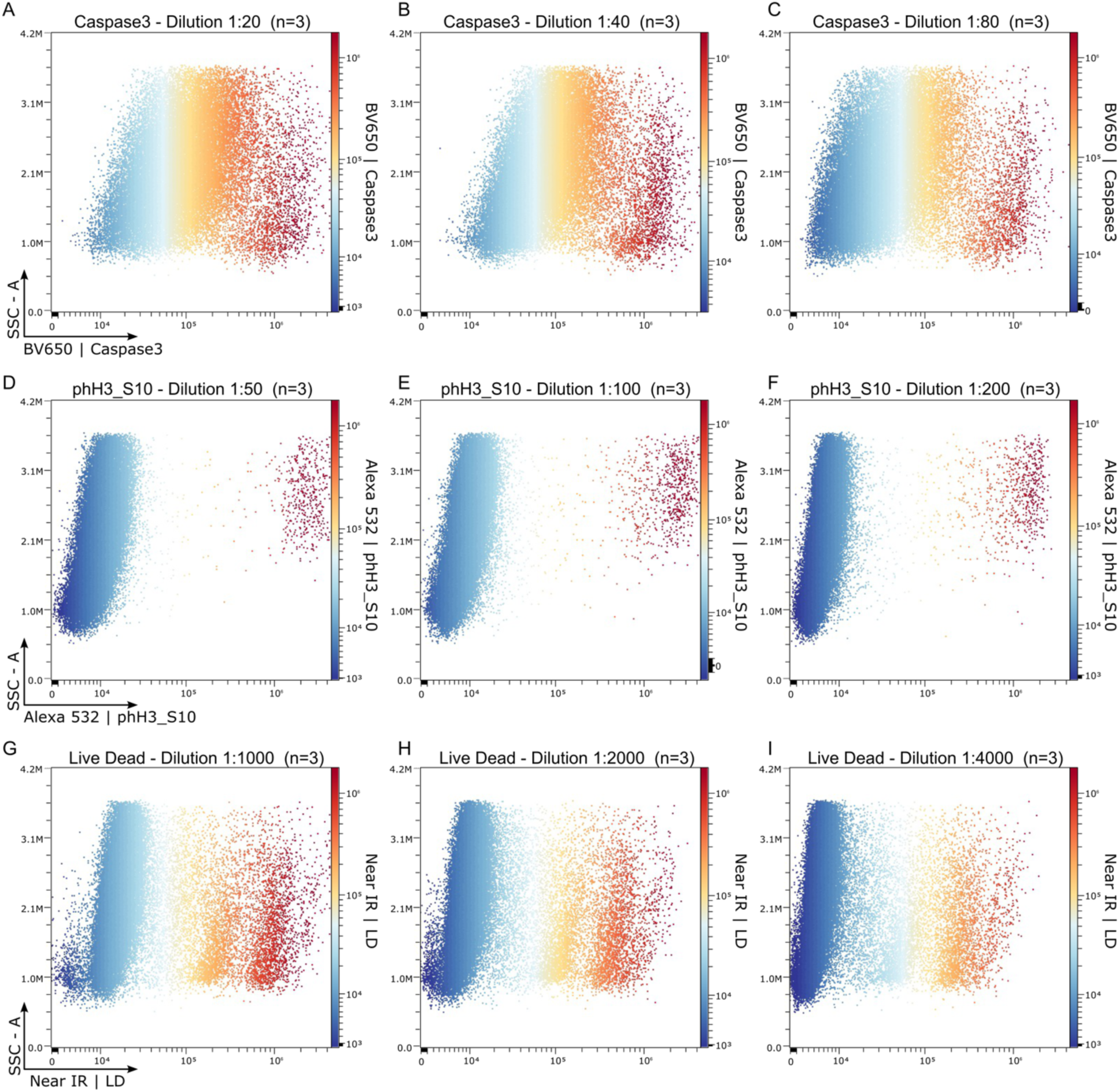
Titration experiment for specific reagents. A-C. Caspase3 antibody was titrated at 1:20, 1;40, and 1:80 dilutions. The 1:200 dilution was selected for experiments. D-F: Phospho-Histone H3 (Ser10) was titrated at 1:50, 1:100, and 1:200. A dilution of 1:400 was chosen for future experiments because the signal was still nearing the detection limits at the highest tested dilution. G-I. Live dead dye staining was titrated at 1:1,000, 1:2,000, and 1:4,000. Similar to phH3, a dilution of 1:8,000 for the Live/Dead stain was selected because the signal was still nearing the detection limits at the highest tested dilution.

