## Supplementary figures and images for "Nuclear Histone 3 Post-Translational Modification Profiling in Whole Cells using Spectral Flow Cytometry"

### Supplementary File 2

**Supplementary File 2.** NxN plots to assess unmixing quality.

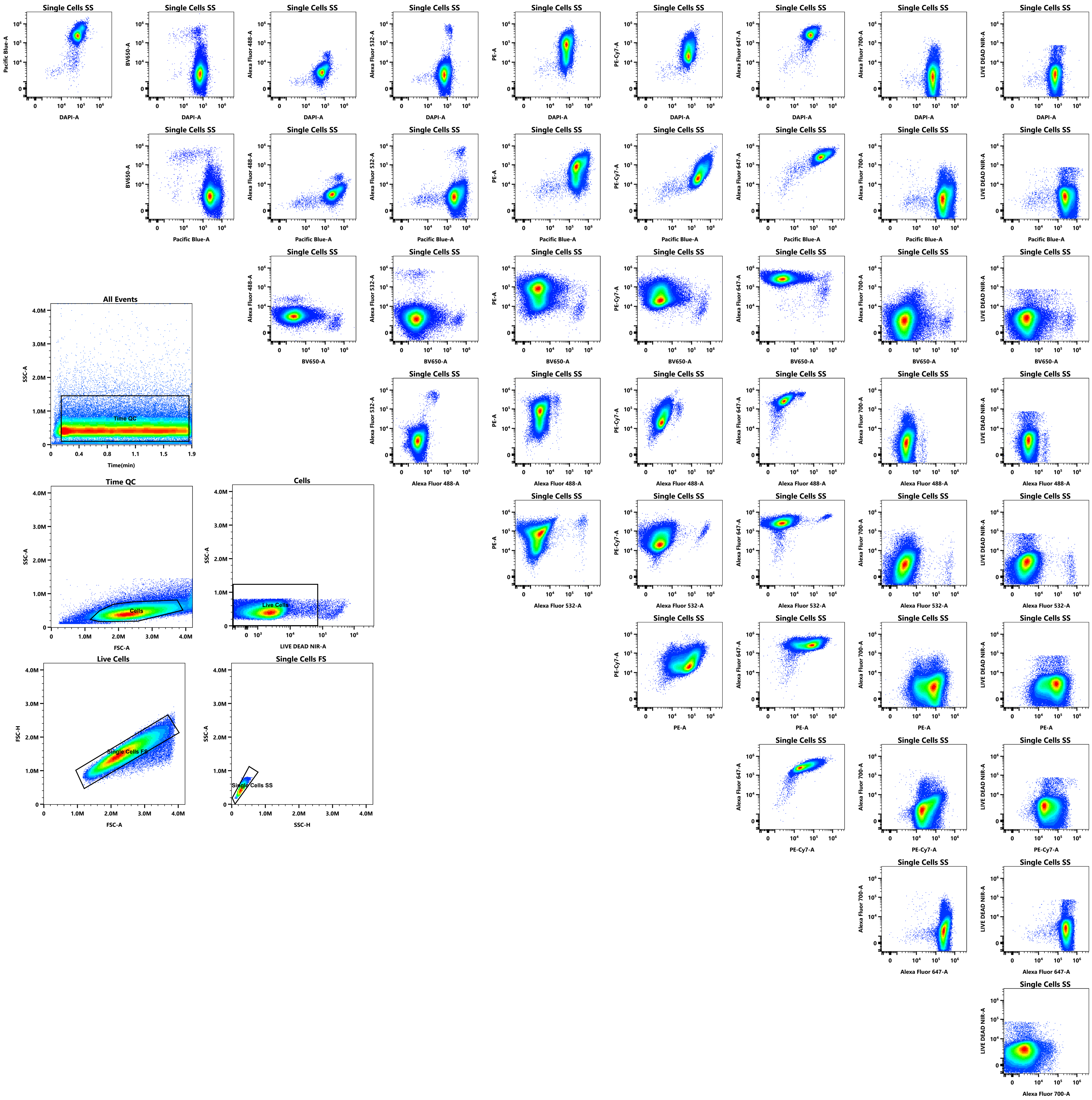
