## Supplementary material for "Nuclear Histone 3 Post-Translational Modification Profiling in Whole Cells using Spectral Flow Cytometry": Table S1

**Table S1. Spectral unmixing controls**

| Laser | Fluorochrome | Marker | Clone | Vendor | Catalog # | Unmixing |
|---|---|---|---|---|---|---|
| 355 | Violet | FxCycle/DNA | NA | Thermo Fisher | R37166 | Cells |
| 405 | Pacific Blue | H3K9ac | C5B11 | Cell Signaling | 11857S | Beads |
| 405 | BV650 | Active Caspase-3 | C92-605.rMAb | BD | 570179 | Beads |
| 405 | mFluorViolet 500 SE | PAX6(1) | PAX6/1166 | Novus Biologicals | 47915MFV500 | Beads |
| 488 | Alexa Fluor 488 | H3K14ac | D4B9 | Cell Signaling | 26828 | Beads |
| 488 | Alexa Fluor 532 | PhosphoH3 (Ser10) | D2C8 | Cell Signaling | 52725 | Beads |
| 561 | PE | PAX6(2) | O18-1330 | BD | 561552 | Beads |
| 561 | PE | H3K4me1(1) | D1A9 | Cell Signaling | 55800S | Beads |
| 561 | PE-Cy7 | H3K27ac | D5E4 | Cell Signaling | 30342 | Beads |
| 640 | Alexa Fluor 647 | H3K4me3(2) | C42D8 | Cell Signaling | 12064 | Beads |
| 640 | Alexa Fluor 700 | Total H3 | D1H2 | Cell Signaling | 54826 | Beads |
| 640 | Near IR 780 | Fixable Viability | NA | BD | 565388 | Cells |

(1) Included in the panel for drug treatment validation assay **(Figure S2)**

(2) Excluded in the panel for drug treatment validation assay **(Figure S2)**
