## Supplementary material for "Nuclear Histone 3 Post-Translational Modification Profiling in Whole Cells using Spectral Flow Cytometry": Table S2

**Reagents**

- Acetyl-Histone H3 (Lys9) (Cell Signaling Technologies, catalog number: 28036S)
- Acetyl-Histone H3 (Lys27) (Cell Signaling Technologies, catalog number: 15562S)
- Histone H3 647 (Cell Signaling Technologies, catalog number: 12230S)
- Mono-Methyl-Histone H3 (Lys4) (Cell Signaling Technologies, catalog number: 55800S)
- Di-Methyl-Histone H3 (Lys4) (Cell Signaling Technologies, catalog number: 31777S)
- Tri-Methyl-Histone H3 (Lys4) (Cell Signaling Technologies, catalog number: 62255S)

**Table S1: Stratedigm Antibody Panel**

| Antibody | Catalog Number | Dilution | Volume | FACs Buffer | Total volume |
|---|---|---|---|---|---|
| Acetyl-Histone H3 (Lys9)* | 28036S | 1:50 | 12 µL | 588 µL | 600 µL |
| Histone H3 647* | 12230S | 1:50 | 12 µL | 588 µL | 600 µL |
| Acetyl-Histone H3 (Lys27)* | 15562S | 1:50 | 12 µL | 588 µL | 600 µL |
| Mono-Methyl-Histone H3 (Lys4)* | 55800S | 1:50 | 12 µL | 588 µL | 600 µL |
| Di-Methyl-Histone H3 (Lys4)* | 31777S | 1:50 | 12 µL | 588 µL | 600 µL |
| Tri-Methyl-Histone H3 (Lys4)* | 62255S | 1:50 | 12 µL | 588 µL | 600 µL |
| Acetyl-Histone H3 (Lys9) **SCC** | 28036S | 1:50 | 6 µL | 294 µL | 300 µL |
| Acetyl-Histone H3 (Lys27) **SCC** | 15562S | 1:50 | 6 µL | 294 µL | 300 µL |
| Mono-Methyl-HistoneH3(Lys4)**SCC** | 55800S | 1:50 | 6 µL | 294 µL | 300 µL |
| Di-Methyl-Histone H3 (Lys4) **SCC** | 31777S | 1:50 | 6 µL | 294 µL | 300 µL |
| Tri-Methy-Histone (Lys4) **SCC** | 62255S | 1:50 | 6 µL | 294 µL | 300 µL |
| Histone H3 647 **SCC** | 12230S | 1:50 | 6 µL | 294 µL | 300 µL |
| Live/Dead **SCC** | L34963 | n/a | n/a | n/a | n/a |

*** Denotes that the antibody can be combined into their respective sample.**
